# Brilacidin, a COVID-19 Drug Candidate, Exhibits Potent *In Vitro* Antiviral Activity Against SARS-CoV-2

**DOI:** 10.1101/2020.10.29.352450

**Authors:** Allison Bakovic, Kenneth Risner, Nishank Bhalla, Farhang Alem, Theresa L. Chang, Warren Weston, Jane A. Harness, Aarthi Narayanan

## Abstract

**Summary:** Severe Acute Respiratory Syndrome Coronavirus 2 (SARS-CoV-2), the newly emergent causative agent of coronavirus disease-19 (COVID-19), has resulted in more than one million deaths worldwide since it was first detected in 2019. There is a critical global need for therapeutic intervention strategies that can be deployed to safely treat COVID-19 disease and reduce associated morbidity and mortality. Increasing evidence shows that both natural and synthetic antimicrobial peptides (AMPs), also referred to as Host Defense Proteins/Peptides (HDPs), can inhibit SARS-CoV-2, paving the way for the potential clinical use of these molecules as therapeutic options. In this manuscript, we describe the potent antiviral activity exerted by brilacidin—a *de novo* designed synthetic small molecule that captures the biological properties of HDPs—on SARS-CoV-2 in a human lung cell line (Calu-3) and a monkey cell line (Vero). These data suggest that SARS-CoV-2 inhibition in these cell culture models is primarily a result of the impact of brilacidin on viral entry and its disruption of viral integrity. Brilacidin has demonstrated synergistic antiviral activity when combined with remdesivir. Collectively, our data demonstrate that brilacidin exerts potent inhibition of SARS-CoV-2 and thus supports brilacidin as a promising COVID-19 drug candidate.

**Highlights:**

- Brilacidin potently inhibits SARS-CoV-2 in an ACE2 positive human lung cell line.
- Brilacidin achieved a high Selectivity Index of 426 (CC_50_=241μM/IC_50_=0.565μM).
- Brilacidin’s main mechanism appears to disrupt viral integrity and impact viral entry.
- Brilacidin and remdesivir exhibit excellent synergistic activity against SARS-CoV-2.

**Significance Statement:** SARS-CoV-2, the emergent novel coronavirus, has led to the current global COVID-19 pandemic, characterized by extreme contagiousness and high mortality rates. There is an urgent need for effective therapeutic strategies to safely and effectively treat SARS-CoV-2 infection. We demonstrate that brilacidin, a synthetic small molecule with peptide-like properties, is capable of exerting potent *in vitro* antiviral activity against SARS-CoV-2, both as a standalone treatment and in combination with remdesivir, which is currently the only FDA-approved drug for the treatment of COVID-19.

## Introduction

The global COVID-19 pandemic resulting from infection by SARS-CoV-2, the novel coronavirus, has resulted in more than 1.16 million deaths worldwide, including 8.7 million cases and over 226,000 fatalities in the United States. **^1^** Additional data indicate there is a significant underreporting of deaths.**^2^** Moreover, the U.S. Congressional Budget Office estimates the pandemic will cost the U.S. economy nearly $8 trillion through 2030,**^3^** with recent analysis estimating aggregate economic losses due to COVID-19 may be as high as $16 trillion.**^4^**

Approximately 15 percent of COVID-19 patients will develop lung injury, including severe respiratory distress that can progress to Acute Respiratory Distress Syndrome (ARDS), often requiring prolonged ventilator support and leading to death.**^5^** Intensive care units, hospitals and health care systems risk becoming overwhelmed by critically ill COVID-19 patients. COVID-19 itself is characterized by a heightened inflammatory component, with several pro-inflammatory cytokines, such as TNF-α, IL-1β, and IL-6, strongly upregulated in infected individuals.**^6, 7, 8^** The prevalence of secondary bacterial infections,**^9, 10, 11, 12^** which can occur in up to 20 percent of cases among hospitalized patients, reinforces the need for a multi-pronged treatment approach that can address the complexities of COVID-19.

To date, no vaccines have been approved to prevent SARS-CoV-2 infection, alongside few modestly effective therapies to treat COVID-19, with remdesivir the only FDA-approved treatment.**^13^** Therapies are predominately being developed to treat acute viral infections in hospital settings versus being evaluated for their prophylactic potential.**^14, 15, 16^** It is likely multiple interventional strategies will be required to prevent and treat COVID-19,**^17, 18^** with SARS-CoV-2 possibly becoming an endemic viral infection. **^19^** Drugs shown to exhibit broad spectrum antiviral activity might help address COVID-19 and potential future viral pandemics.**^20^**

Natural and synthetic antimicrobial peptides (AMPs), also called Host Defense Proteins/Peptides (HDPs), comprise potentially effective countermeasures against COVID-19, having shown inhibitory activity against multiple viruses. **^21, 22, 23, 24, 25, 26, 27, 28, 29, 30^** Integral components of the innate immune response, HDPs are typically small (12-80 amino acids) proteins and peptides expressed widely in the animal kingdom that serve as the “first line of defense” against foreign pathogens and potential subsequent infection and related inflammation. HDPs have been recognized as potential sources for promising therapeutics. **^31, 32, 33, 34, 35, 36, 37, 38, 39^** In mammals, HDPs are found within granules of neutrophils and in secretions from epithelial cells covering skin and mucosal surfaces. Their discovery can be traced to 1939, with the extraction of gramcidin from *Bacillus brevis*, followed, in the 1980s, by the isolation of cecropins in silk moths (Hyalophora) by Hans Boman**^40^** and magainins in frogs (*Xenopus laevis*) by Michael Zasloff.**^41^**

Despite a variety of chemical sequences, as well as secondary and tertiary structures, most HDPs share an amphiphilic topology—with a positively-charged face on one side and a hydrophobic face on the other side.**^42^** In contrast, most bacteria (and some viruses) express a negative charge on their outer membranes, **^43^** as well as lack cholesterol, an essential component of mammalian membranes.**^44, 45^** Due to these differences, HDPs can selectively and variously interact with pathogens, increasing their susceptibility to proteolysis and degradation. **^46^** The immunomodulatory activity of HDPs is believed to be underestimated since most HDP research has been conducted in cell cultures and not observed directly in humans.**^47, 48^** It has been suggested that clinical development efforts against COVID-19 should be focused on advancing and evaluating drugs with immunomodulating properties to trigger the adaptive immune response and control inflammation.**^49^** The U.S. National Institutes of Health has announced a large Phase 3 clinical trial to test immune modulators against COVID-19.**^50^** Other modes of action for HDPs, including an ability to affect multiple targets at once, continue to be explored, as this attribute may be integral to contributing to overall HDP efficacy.**^51^**

SARS-CoV-2 infection has been shown to suppress defensins, a type of HDP, suggesting that the innate immunity provided by defensins may be compromised.**^52^** Suppression of LL37, a human cathelicidin (another type of HDP) shown to possess immunomodulatory and antiviral properties,**^53^** has also been observed in COVID-19 patients.**^54^** Increased expression of defensins and cathelicidins, e.g., LL37, can decrease both the viral and inflammatory load in the context of several respiratory viral infections,**^55^** further supporting the protective role of HDPs. LL-37 has even been studied in an exploratory clinical trial in which encouraging early results were reported.**^56^** HDPs, in particular human defensins, have been proposed as adjuvants for vaccine development.**^57, 58, 59^** Due to their unique physico-chemical properties (hydrophobicity, cationic charge) enabling impact to viruses prior to infection, HDPs are well-suited for prophylactic development, as some HDPs appear to behave in a manner similar to the neutralization of virus observed in the presence of neutralizing antibodies.**^60, 61, 62, 63^** Numerous recently published reports**^64, 65, 66, 67, 68, 69, 70, 71, 72^** support the antiviral activity of HDPs in the context of coronaviruses,**^73^** including SARS-CoV-2.

Brilacidin (PMX-30063) is a synthetic, non-peptidic, small molecule mimetic of HDPs.**^74, 75, 76, 77, 78^** Building on “first principles” in medicinal chemistry, rational design tools—leveraging sophisticated informatics to fine-tune physico-chemical properties and structure-activity relationships **^79, 80, 81^** —enabled brilacidin (an arylamide foldamer **^82, 83^**) to overcome the shortcomings and challenges that have complicated the clinical development of natural HDPs.**^84^** These include: proteolytic degradation, toxicity, lack of efficacy, malabsorption, and high cost to produce. In contrast to natural HDPs, as well as other HDP analogs, brilacidin was designed *de novo***^85, 86, 87^** to be much smaller, more stable, more potent, more selective, and more economical to manufacture.

Within this broader context of the global COVID-19 pandemic and the potential therapeutic role for HDPs, brilacidin was evaluated in laboratory testing to determine if the drug might exhibit antiviral properties against SARS-CoV-2.

## Results

### Brilacidin inhibits SARS-CoV-2 replication (Vero Cells)

As a first step, the potential of brilacidin to exert an antiviral activity against SARS-CoV-2 was assessed using Vero cells as an infection model. Toxicity assessment of brilacidin in Vero cells was initially performed by incubating the cells with increasing concentrations of the compound for 24 hours after which cell viability was assessed by Cell Titer Glo assay. Brilacidin at up to 40μM concentration did not affect cell viability when compared to the DMSO vehicle control; a dose-dependent, statistically significant decrease in cell viability was detected at higher concentrations. The effect of brilacidin treatment on SARS-CoV-2 viral replication was then evaluated in Vero cells by plaque assay. Vero cells were pre-treated with brilacidin for 2 hours after which media containing the drug was removed and replaced with virus inoculum (Washington strain 2019-nCoV/USA-WA1/2020). Infection was allowed to progress for one hour after which the inoculum was removed and replaced with brilacidin containing media. Cell culture supernatants from vehicle-treated and brilacidin-treated cells were collected at 16 hours post infection, and the SARS-CoV-2 infectious titer in the supernatants quantitated by plaque assay and compared to the DMSO-treated control. The data demonstrate that brilacidin treatment resulted in a dose-dependent decrease in infectious viral titer with a maximum of 53% inhibition of virus observed in the presence of the higher concentration of compound (10μM) that was tested (**Figure 2A**).

**Figure 1.**
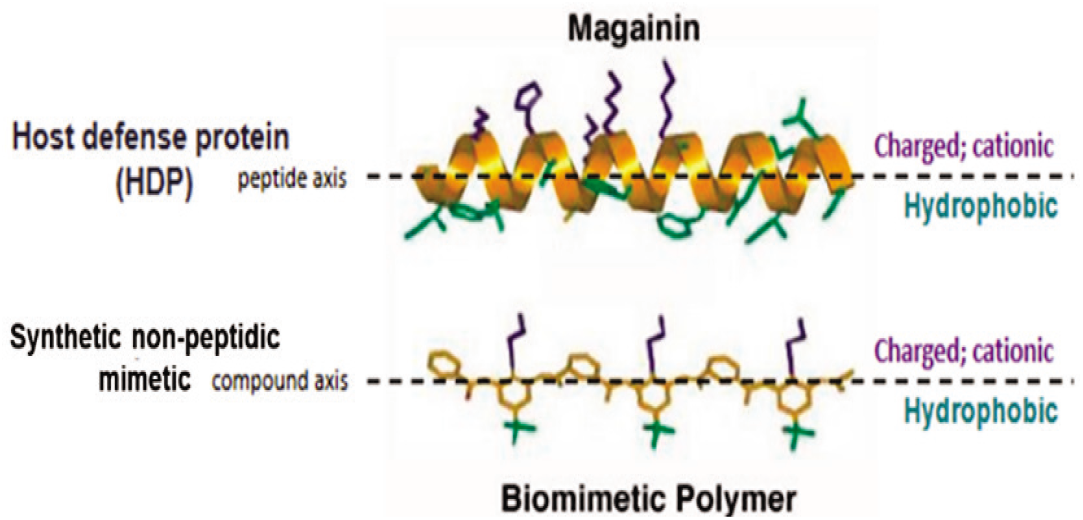
Structure of Brilacidin. Schematic representation of the amphiphilic α-helix structure of the HDP magainin above that of a synthetic non-peptidic mimetic polymer, such as brilacidin, capturing the structural and biological properties of HDPs using fully synthetic, non-peptidic scaffolds and sidechains.

**Figure 2A.**
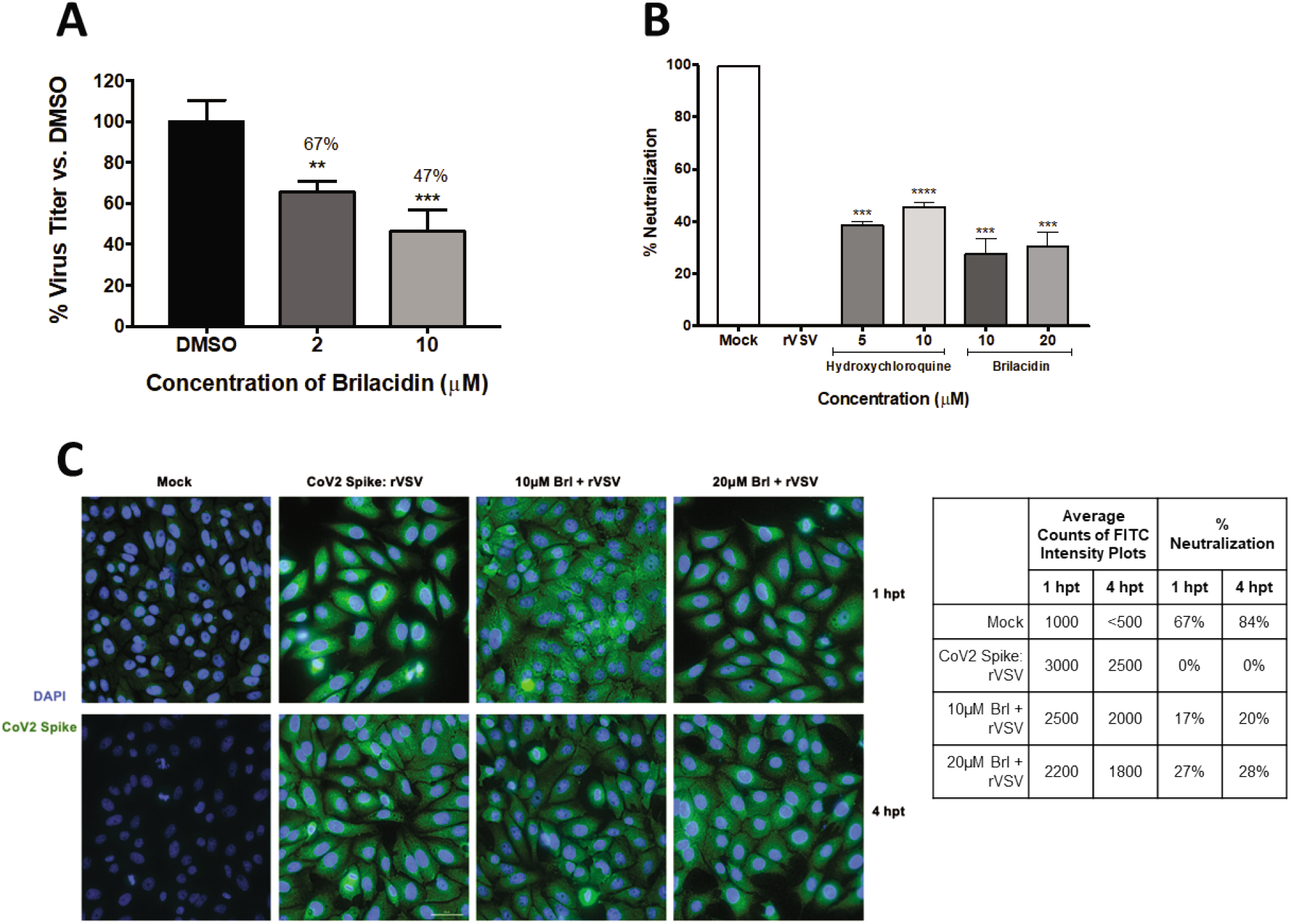
Brilacidin inhibits SARS-CoV-2 replication (Vero cells) (A) Vero cells were pretreated for 2h with 2 or 10μM brilacidin, infected with SARS-CoV-2 non-directly at MOI 0.1 for 1h, and post-treated with media containing brilacidin as described in Materials and Methods. At 16hpi, viral supernatants were evaluated by plaque assay as described in Materials and Methods. **Figure 2B, 2C. Brilacidin appears to impact entry of SARS-CoV-2 (Vero cells)** (B) Brilacidin was measured at 10 and 20μM for neutralization activity against a luciferase-expressing pseudotyped virus (rVSV) containing the SARS-CoV-2 spike protein using luciferase assay in Vero cells at 24hpt as described in Materials and Methods and compared to neutralization activity of hydroxychloroquine. (C) Vero cells were treated with 10 or 20μM brilacidin for neutralization activity against SARS-CoV-2 rVSV, and cells imaged and quantified using fluorescent microscopy and FITC surface intensity plots at 1 and 4hpt as described in Materials and Methods. Graphs are representative of one independent experiment performed in technical triplicates (n=3). Brl indicates brilacidin. **p<0.0021, ***p<0.0002, ****p<0.0001.

### Brilacidin appears to impact entry of SARS-CoV-2 (Vero cells)

The potential for brilacidin to interfere with viral entry was assessed in the Vero cell line by looking at the ACE2:Spike protein interaction, using a rVSV pseudotyped SARS-CoV-2 expressing a luciferase reporter gene. The pseudovirus retains the SARS-CoV-2 spike protein on its surface and is capable of ACE2 based viral entry, which can be quantitated by measuring intracellular luciferase expression; the pseudovirus is not capable of viral RNA synthesis once inside the cell, hence any inhibitory effect is regarded as most likely limited to the early entry and post-entry steps. The inhibition of pseudovirus (rVSV) attachment and entry into cells was quantified in the context of brilacidin treatment (10μM and 20μM), and hydroxychloroquine was utilized as a control. The data demonstrate that brilacidin treatment inhibited the pseudovirus at both concentrations tested in a comparable manner (**Figure 2B**). The inhibition observed in the context of the replication-incompetent pseudovirus was supportive of the suggested role of brilacidin as an inhibitor of viral entry and potential early post-entry steps. To further support this observation, confocal microscopy was performed using an antibody directed against the viral spike protein in the presence and absence of brilacidin using the pseudovirus (**Figure 2C**). Quantification of the confocal images revealed that incubation of SARS-CoV-2 with brilacidin resulted in a decreased intracellular spike protein signal at 1 and 4 hours post infection.

### Brilacidin appears to disrupt the integrity of the SARS-CoV-2 virion

The potential for brilacidin to interfere directly with the intact virus prior to cell attachment was assessed. If brilacidin is able to impact viral integrity, inhibition of SARS-CoV-2 (Washington strain) should increase above that observed in the assay (as in **Figure 2A**) when modified to include a brilacidin-treated inoculum. To evaluate this, the inoculum was independently incubated with 10μM brilacidin for 1 hour after which the treated inoculum (virus-brilacidin mix) was used to infect Vero cells. The infection was, thus, also carried out in the presence of brilacidin for one hour, after which the inoculum was removed and replaced with media containing the drug. The culture supernatants were assessed for viral load by plaque assay at 24 hours post infection. The outcomes of this experiment revealed a higher inhibition of SARS-CoV-2 (72% inhibition), alluding to an inhibitory activity exerted upon the virus directly (**Figure 2D**). Using the same assay (for 10μM brilacidin), the intracellular viral genomic copy numbers were assessed by semi-quantitative RT-PCR at 24 hours post infection, which demonstrated a 29% decrease in the viral genomic copies with brilacidin treatment; this extent of inhibition of intracellular RNA copies assessed at later timepoints in infection is not unexpected for an inhibitor with likely activity exerted during the early entry and post-entry steps.

**Figure 2D, 2E.**
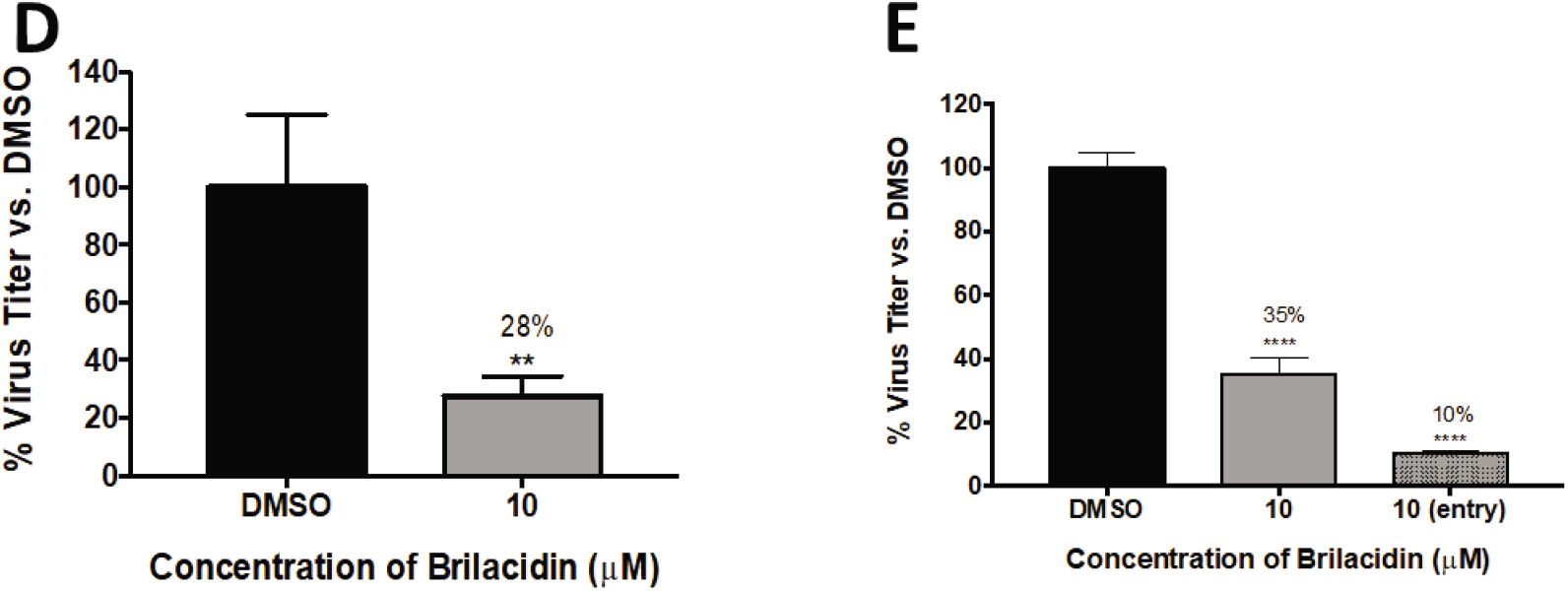
Brilacidin appears to disrupt the integrity of the SARS-CoV-2 virion. (D) Vero cells were pre-treated for 2h with 10μM brilacidin. SARS-CoV-2 was diluted to MOI 0.1 in culture media containing brilacidin and incubated for 1h. Viral inoculum containing inhibitor was added to cells for 1h for a direct infection, and post-treated with media containing brilacidin as described in Materials and Methods. At 24 hpi, viral supernatants were evaluated by plaque assay as described in Materials and Methods. (E) Vero cells were pretreated for 2h with 10μM brilacidin or media only (indicated as [entry]). SARS-CoV-2 was diluted to MOI 0.1 in culture media containing 10μM brilacidin and incubated for 1h. Brilacidin-treated viral inoculum was added to cells for 1h for a direct infection, and replaced with inhibitor containing media or media only (indicated as [entry]) as described in Materials and Methods. At 24hpi, viral supernatants were evaluated by plaque assay as described in Materials and Methods. Graphs are representative of one independent experiment performed in technical triplicates (n=3). **p<0.0021, ****p<0.0001.

To independently assess the impact of brilacidin on the virion and thus add support to the role of brilacidin as an inhibitor of viral entry, a direct virus inhibition assay was conducted akin to virus neutralization observed in the presence of antibodies. To that end, SARS-CoV-2 inoculum was incubated with brilacidin at 10μM concentration for 1 hour after which the treated inoculum was used to infect Vero cells. In this experiment, the cells were not pre-treated with the inhibitor (brilacidin) prior to the infection. After 1 hour of infection, the inoculum was removed and replaced with fresh media without any inhibitor and the cells were maintained in inhibitor-free media for 24 hours. The infectious virus titer in the supernatant was quantified by plaque assay, which revealed a dramatic 90% reduction of virus titer (**Figure 2E**, indicated as [entry]). This inhibition was approximately 25% higher than that observed when the cells were pre- and post-treated with brilacidin concomitantly (**Figure 2E**) supporting the concept that brilacidin has a direct inhibitory effect on the virus in a manner similar to the neutralization of antibodies, potentially by disrupting viral integrity and thus impairing the virion’s ability to complete the viral entry process.

### Brilacidin exhibits potent inhibition of SARS-CoV-2 in a human cell line (Calu-3 cells)

To ascertain that brilacidin can elicit anti-SARS-CoV-2 activity in an ACE2 positive human lung cell, experiments were conducted in the Calu-3 infection model. The toxicity of brilacidin in this cell line was initially assessed at 10μM and 20μM concentrations by incubating the cells with the compound for 24 hours. The assay revealed that these concentrations of brilacidin were nontoxic to Calu-3 cells. The inhibitory effect of brilacidin in the Calu-3 cell line was first confirmed by pre-treatment (for 2 hours) and post-infection treatment (for 24 hours) of cells with brilacidin, which demonstrated a dose-dependent decrease of viral load, with the higher concentration of brilacidin providing 61% inhibition of infectious viral titer (**Figure 3A**). However, when the experiment was modified to include a brilacidin-treated inoculum – with direct pre-incubation of the virus with brilacidin for 1 hour prior to infection, and with infection carried out in the presence of the compound – the extent of inhibition dramatically increased, resulting in 95% and 97% reduction of infectious viral titer at the 10μM and 20μM concentration of the compound respectively (**Figure 3B**). Quantification of intracellular viral RNA by semi-quantitative RT-PCR at 24 hours post infection (for 10μM brilacidin) demonstrated a 33% decrease in the viral genomic copies upon brilacidin treatment.

**Figure 3.**
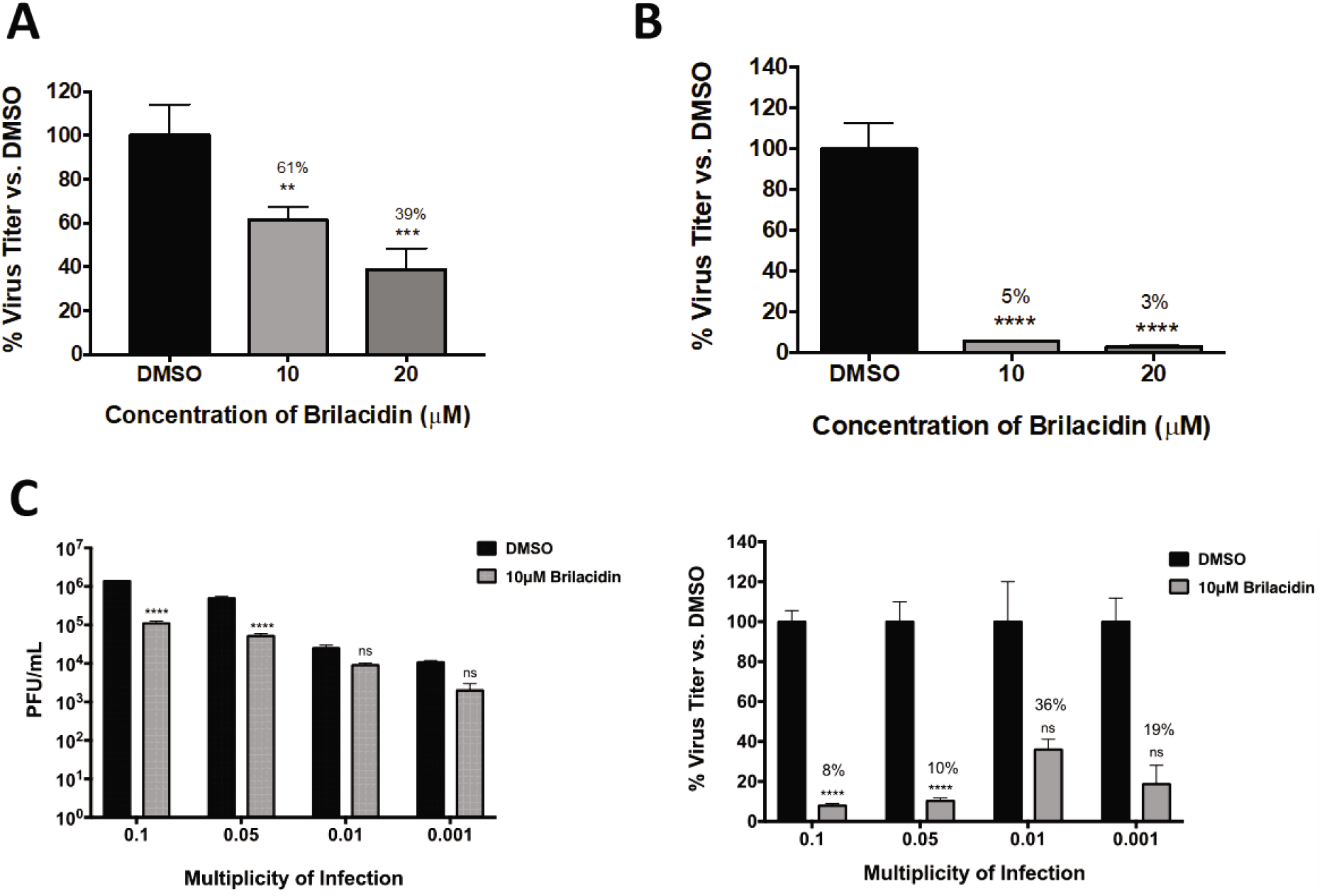
Brilacidin exhibits potent inhibition of SARS-CoV-2 in a human cell line (Calu-3 cells) (A) Calu-3 cells were pretreated for 2h with 10 or 20μM brilacidin and infected with SARS-CoV-2 at MOI 0.1 non-directly or (B) directly with brilacidin for 1h as described in Materials and Methods. Cells were post-treated with media containing brilacidin, and at 24hpi viral supernatants were evaluated by plaque assay as described in Materials and Methods. (C) Calu-3 cells were pretreated for 2h with 10μM of brilacidin, infected with SARS-CoV-2 at MOI of 0.1, 0.05, 0.01, or 0.001 directly with brilacidin at 10μM for 1h, and post-treated with media containing brilacidin as described in Materials and Methods. At 24hpi, viral supernatants were evaluated by plaque assay as described in Materials and Methods. Statistical analyses for varying MOI was determined using Two-Way ANOVA with Sidak’s multiple comparisons test. Significance for all other graphs were determined as described in Materials and Methods. Graphs are representative of one independent experiment performed in technical triplicates (n=3). **p<0.0021, ***p<0.0002, ****p<0.0001, ns = not significant.

The inhibition of infectious virus titer as a variable of viral load was assessed by quantifying inhibition at lower multiplicities of infection (MOI) with brilacidin at a fixed concentration of 10μM. Interestingly, the inhibitory potential of brilacidin was best observed at the highest MOIs tested, with inhibition of virus at the lower MOIs (0.01 and 0.001) not showing statistical significance (**Figure 3C**). The inhibition exerted at the MOIs of 0.1 and 0.05 were extremely comparable to each other.

### Selectivitiy Index determination for brilacidin against SARS-CoV-2 in a human cell line (Calu-3 cells)

The Selectivity Index, a ratio that compares a drug’s cytotoxicity and antiviral activity, is a measure of how likely a drug is to be safe and effective when translated to human testing in the clinic. The 50% cytotoxicity concentration (CC_50_), i.e., the concentration that results in the reduction of cell viability by 50%, is compared to the concentration thar results in 50% of the maximal inhibitory response (IC_50_). The values for 90% cell viability (CC_10_) and the 90% inhibitory concentration (IC_90_) were also derived. The CC_50_ values for brilacidin in the context of Calu-3 cells were assessed by measuring cell survival over a concentration range between 0.1 – 200μM, which revealed that the 50% reduction in cell viability was observed at a concentration of 241μM, with 90% viability (CC_10_) observed at 26.8μM, thus suggesting brilacidin was extremely well tolerated (**Figure 4A**). Quantification of the inhibitory response – when the virus was directly pre-incubated with brilacidin prior to infection; cells were treated prior to infection; brilacidin was present during infection; and infected cells were maintained in the presence of brilacidin post-infection (assay as in **Figure 3B**) – demonstrated brilacidin achieved 90% inhibition at a concentration of 2.63μM and 50% inhibition at 0.565μM, yielding a Selectivity Index of 426 (CC_50_=241μM/IC_50_=0.565μM). (**Figure 4B**)

**Figure 4.**
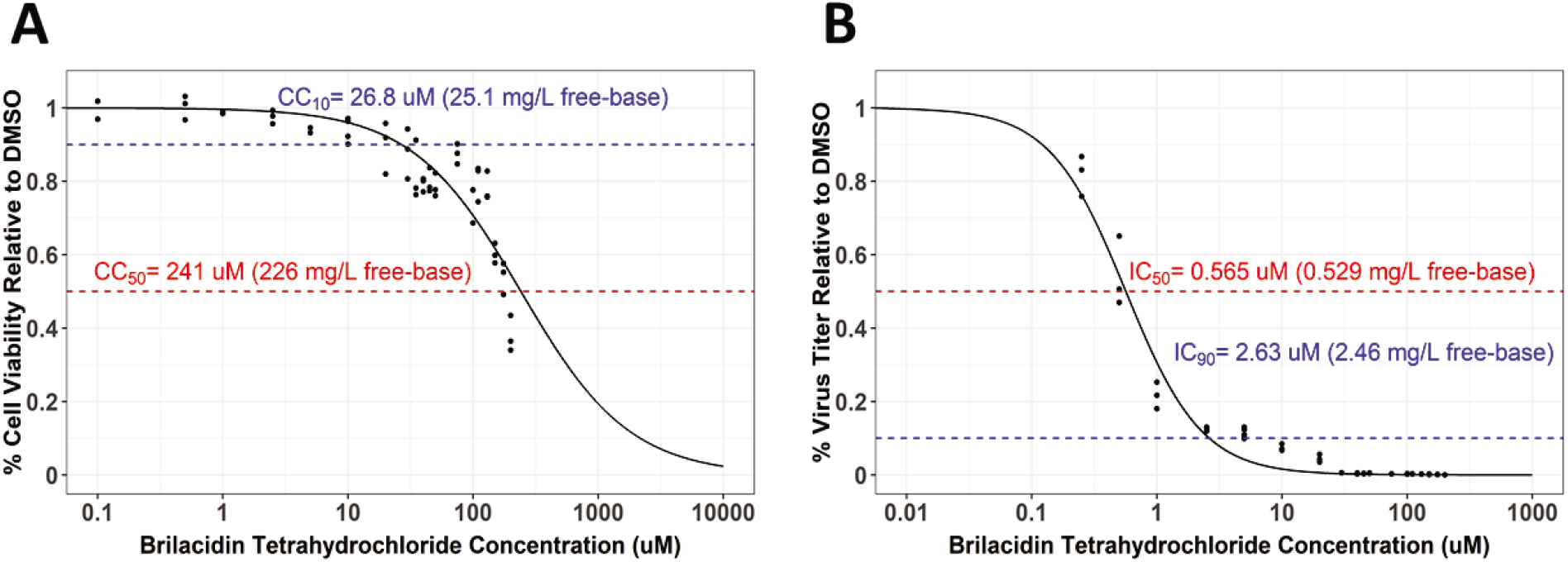
Brilacidin exhibits potent inhibition of SARS-CoV-2 in a human cell line (Calu-3 cells) (A) Calu-3 cells were treated at increasing concentrations of brilacidin at a range of 0.1-200μM. Cell viability was measured at 24hpt and calculated versus the DMSO control as described in Materials and Methods. (B) Calu-3 cells were pre-treated for 2h with brilacidin at increasing concentrations, directly infected with treated (and pre-incubated) viral inoculum at MOI 0.1 at the indicated pre-treatment brilacidin concentration for 1h, and post-treated with media containing brilacidin as described in Materials and Methods. At 24hpi, viral supernatants were evaluated by plaque assay as described in Materials and Methods. Graphs are of one independent experiment performed in technical triplicates (n=3). Sigmoidal Hill-type models as a function of brilacidin tetrahydrochloride concentration were fit to the cell viability (A) and inhibitory response (B) data using non-linear least squares regression. The dashed lines indicate (A) derived CC_10_ and CC_50_ values, and (B) derived IC_50_ and IC_90_ values. The calculated Selectivity index is 426 (CC_50_=241μM/IC_50_=0.565μM).

### Brilacidin in combination with other antiviral treatments; synergistic activity against SARS-CoV-2 in combination with remdesivir (Calu-3 cells)

As brilacidin appears to act primarily by disrupting viral integrity and blocking viral entry, combining the drug with antiviral treatments that have a different mechanism of action may result in synergistic inhibition when administered in combination. The potential for brilacidin to exert a synergistic inhibition of SARS-CoV-2 when combined with current frontline COVID-19 antiviral treatments, namely, remdesivir and favipiravir, was assessed. Potential toxicity of combinations of remdesivir or favipiravir with brilacidin were initially assessed in the Calu-3 cell line at 24 hours post treatment. No apparent toxicity could be detected up to 10μM concentration of each of the drugs in the combination regimen. To evaluate the efficacy of combining remdesivir or favipiravir with brilacidin, the cells were pre-treated with brilacidin for 2 hours. The virus inoculum was also independently pre-incubated with brilacidin for 1 hour, and then the treated inoculum was overlaid on cells and the infection allowed to proceed for 1 hour in the presence of brilacidin. Post-infection, the inoculum was removed and media containing both brilacidin and remdesivir or favipiravir or each drug alone for efficacy comparison, was added to the infected cells. Supernatants were obtained at 24 hours post infection and infectious titer quantified by plaque assay. The data revealed that brilacidin and favipiravir independently exerted up to 90% and 80% inhibition respectively and the extent of inhibition did not increase over that exerted by brilacidin alone when the two drugs were used in combination (**Figure 5A**). In contrast, combination of brilacidin with remdesivir at 10μM and 2.5μM concentrations, respectively, reduced the viral infectious titer by >99%, thus providing a highly effective inhibition profile (**Figure 5B**) and achieving greater inhibition than with either compound alone. This synergistic inhibition continued to remain higher than 99% when the concentrations of both compounds were equal (2.5μM each) (**Figure 5C**).

**Figure 5A.**
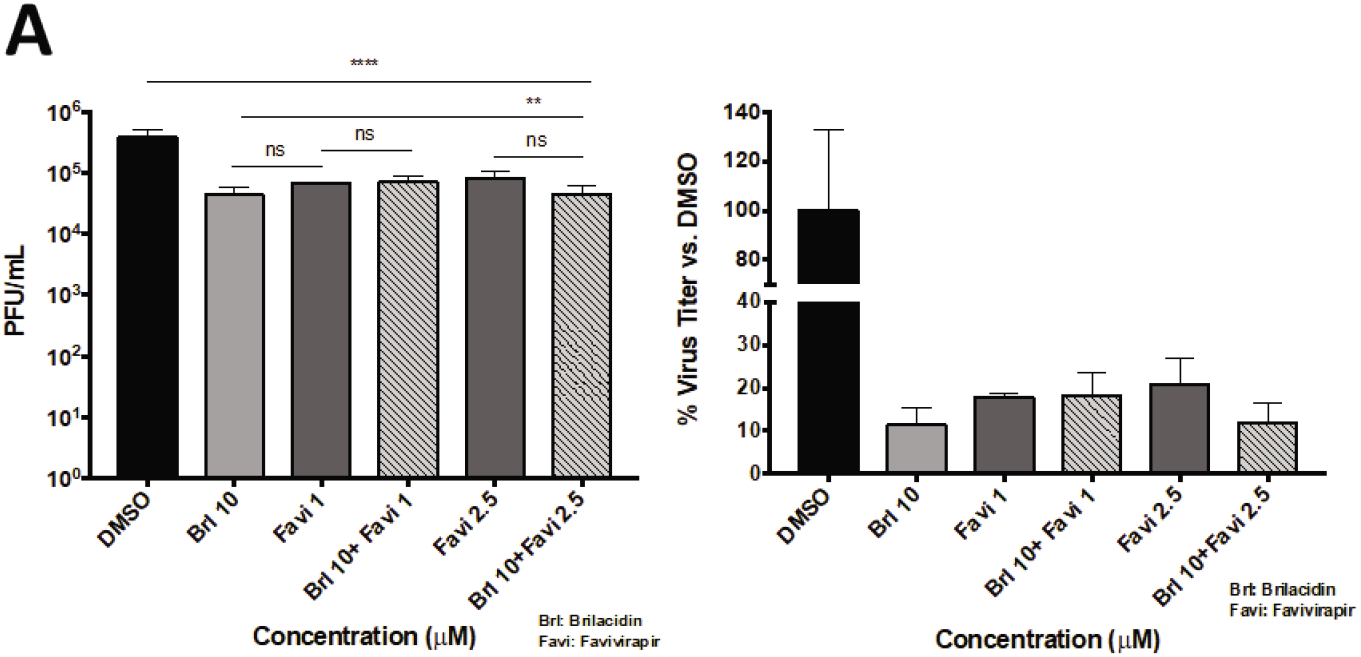
Brilacidin against SARS-CoV-2 in combination with favipiravir (Calu-3 cells) (A) Calu-3 cells were pretreated for 2h with media alone or media containing brilacidin at 10μM (for synergy treatments). Cells were infected with SARS-CoV-2 at MOI 0.05 directly with brilacidin at 10μM (for synergy treatments) or SARS-CoV-2 incubated in media alone (for favipiravir treatment alone). After 1h, post-treatment with favipiravir (B) alone, or mixed with 10μM brilacidin, were added to cells at 1 or 2.5μM concentrations. At 24hpi, viral supernatants were evaluated by plaque assay as described in Materials and Methods. Graphs are representative of one independent experiment performed in technical triplicates (n=3). Brl indicates brilacidin, Favi indicates favipiravir. Statistical analyses for synergy vs. individual control treatments was determined using Unpaired Two-Tailed Student’s t-test. Significance against DMSO was determined as described in Materials and Methods. **p<0.0021, ****p<0.0001, ns = not significant.

**Figure 5B, 5C.**
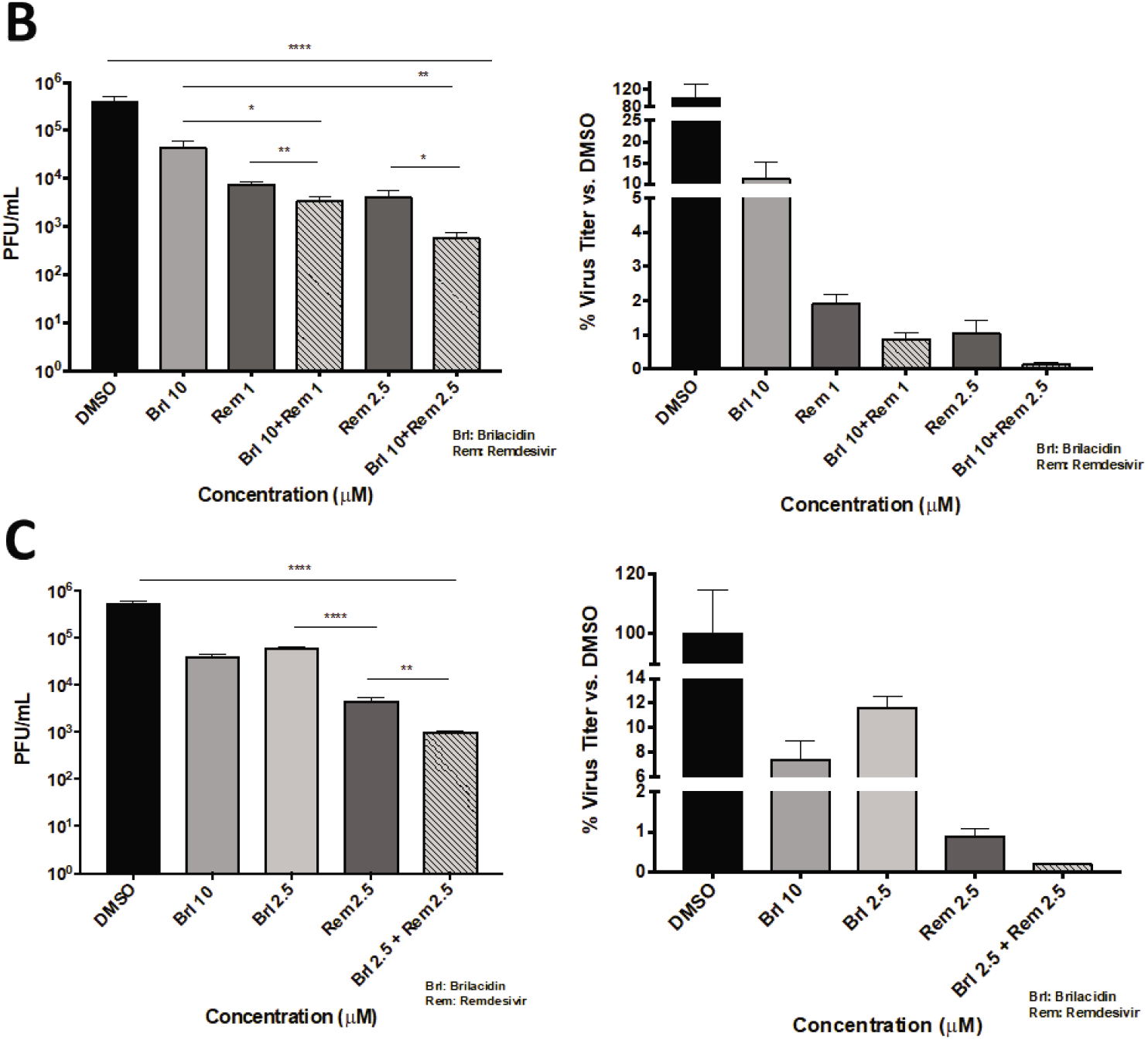
Brilacidin against SARS-CoV-2 in combination with remdesivir (Calu-3 cells) (B) Calu-3 cells were pretreated for 2h with media alone or media containing brilacidin at 2.5 or 10μM (for synergy treatments). Cells were infected with SARS-CoV-2 at MOI 0.05 directly with brilacidin at 2.5 or 10μM (for synergy treatments) or SARS-CoV-2 incubated in media alone (for remdesivir treatment alone). After 1h, post-treatment with remdesivir alone, or mixed with 10μM (B) or 2.5μM (C) brilacidin, were added to cells at 1 or 2.5μM concentrations. At 24hpi, viral supernatants were evaluated by plaque assay as described in Materials and Methods. Graphs are representative of one independent experiment performed in technical triplicates (n=3). Brl indicates brilacidin, Rem indicates remdesivir. Statistical analyses for synergy vs. individual control treatments was determined using Unpaired Two-Tailed Student’s t-test. Significance against DMSO was determined as described in Materials and Methods. *p<0.0332, **p<0.0021, ****p<0.0001.

## Discussion

The ongoing global COVID-19 pandemic powerfully reinforces the need for therapeutic strategies that can safely and effectively address virus- and host-based events elicited during SARS-CoV-2 infection.

In multiple studies, we have attempted to evaluate the capability of brilacidin to decrease viral load in the context of the SARS-CoV-2 infection. Our experiments in the Vero cell line model demonstrate brilacidin decreases viral load in a robust manner when the virus is pre-incubated with brilacidin (**Figure 2D**), suggesting brilacidin impacts the virus directly. This observation was supported by the inhibition seen in the context of a replication incompetent pseudovirus (**Figure 2B, Figure 2C**), further indicating brilacidin’s inhibition during early stages of viral infection. As the pseudovirus expresses the SARS-CoV-2 spike protein on the surface, the data may also support brilacidin’s ability to interfere with the interaction between the viral spike protein and the cellular ACE2 receptor. Brilacidin’s ability to decrease viral load in an ACE2 positive cell line is demonstrated in **Figure 3, Figure 4,** and **Figure 5,** in which Calu-3 cells were used. Additional testing conducted in Caco-2 cells and primary lung fibroblasts obtained from human donors also supported brilacidin’s inhibitory properties in ACE2 positive cell lines (data not shown).

All experiments conducted in Vero and Calu-3 cell line models were supportive of an early inhibition exerted by brilacidin on SARS-CoV-2, indicating the drug’s impact on viral integrity. The idea that brilacidin directly interferes with the integrity of the virion, and hence restricts the virion’s ability to interact with the ACE2 receptor to facilitate the entry process, is further supported by the observation that when drug treatment was limited to the virus alone (**Figure 2E**), with no treatment of host cells, a robust decrease of viral load was still observed. This mechanism of inhibition may be akin to that achieved by neutralizing antibodies that may interact with specific exposed epitopes on the surface of virions. It remains to be determined if the impact of brilacidin on viral membranes is driven by specific viral membrane compositions. Additional studies are planned.

Potent membrane destabilization by brilacidin has been reported in the context of bacteria. Mechanistic-based membrane depolarization studies comparing brilacidin to the lipopeptidic antibiotic daptomycin and the AMP LL16 against *Staphylococcus aureus* showed that brilacidin caused an upregulation of the VraSR and WalKR regulons and affected cytoplasmic proteases and chaperones.**^88^** These findings indicate brilacidin causes significant cell wall stress and additional internal stress due to the accumulation of misfolded proteins. Additional mechanistic studies of brilacidin analogs against *Escherichia coli*, in comparison to the antibiotic polymyxin B, further support this class of compounds ability to destabilize bacterial membranes.**^89^**

While brilacidin’s mechanism of action appears primarily to be extracellular, it may also impact intracellular viral replication and is planned to be researched further. Supportive of this, an *in silico* quantum mechanical molecular screening study of 11,522 compounds identified brilacidin as a potential inhibitor of SARS-CoV-2 based on the potential of its physico-chemical properties to interfere with the intracellular replication of SARS-CoV-2’s main protease (Mpro).^90^

The high CC_50_ (a measure of cytotoxicity) and low IC_50_ (a measure of potency) values observed for brilacidin in Calu-3 cells—yielding a Selectivity Index (SI) for brilacidin of 426 (CC_50_=241μM/IC_50_=0.5 65μM)—strongly support brilacidin’s treatment potential to achieve positive antiviral outcomes in humans. A vast majority of other drugs being evaluated as potential COVID-19 treatments, including repurposed drugs, have SIs that are much lower than that achieved by brilacidin,**^91^** with most drugs failing to show anti-SARS-CoV-2 potency in the <1μM range.**^92^** Of note, the IC_50_ (0.565μM) and IC_90_ (2.63μM) values for brilacidin observed in the Calu-3 cell line are well below clinically-achievable concentrations based on pharmacokinetics observed in a Phase 2b clinical trial of brilacidin for the treatment of Acute Bacterial Skin and Skin Structure Infections (ABSSSI): median Cmax in plasma was 7.67μM brilacidin (free-base) from a single IV dose of 0.6 mg/kg.

A desirable outcome for any potential COVID-19 therapeutic will be its ability to synergize with existing COVID-19 treatments, particularly if the mechanisms of action of the synergistic treatments can impact more than one step of the viral lifecycle. Such combinations are more likely to elicit an additive response, while also reducing the likelihood of viral resistance developing. Along these lines, we conducted experiments to evaluate the potential of brilacidin to work in conjunction with remdesivir and favipiravir (**Figure 5**), two frontline COVID-19 treatments, which proved supportive of synergistic inhibition between brilacidin and remdesivir. Remdesivir and favipiravir are inhibitors that impact the viral RNA synthesis step of the infectious process. These drugs may help decrease progeny viral genomes in infected cells, but they will not be conducive to inhibiting progressive infection of naïve cells once the progeny virions have been released from infected cells.

By combining remdesivir or favipiravir with brilacidin, a two-pronged strategy of inhibiting viral entry and viral RNA synthesis might be successfully leveraged to most effectively control progression of SARS-CoV-2 infection. Furthermore, remdesivir and favipiravir do not possess intrinsic anti-inflammatory activity, unlike brilacidin. Any reduction in inflammatory mediators, including IL-6 and IL-1β in the context of remdesivir and favipiravir treatment, are likely to be a direct consequence of decrease in replicating virus. Moreover, ARDS and associated organ failure observed in the context of COVID-19 are typically later-stage manifestations of disease that are not directly related to, or dependent on, a mounting viral load. Consequently, use of a drug strategy that exhibits intrinsic anti-inflammatory activity will add value to controlling later onset inflammatory damage in COVID-19 patients, potentially occurring beyond the time of active viral multiplication. Along similar lines, it remains to be determined if the combination of convalescent serum—a current COVID-19 treatment that has received FDA Emergency Use Authorization— with brilacidin may confer a protective advantage to patients. It is less likely that a synergistic effect may be observed by combining neutralizing antibodies with brilacidin as it appears the dominant mechanism of action in both cases is disruption of viral entry into cells.

Clearly, an effective COVID-19 therapeutic (or therapeutics in combination) ideally would control both viral load and the corresponding inflammatory damage due to SARS-CoV-2,^93^ and mitigate bacterial co-infections. Exhibiting three-in-one properties—antiviral, immuno/anti-inflammatory, and antibacterial—brilacidin is being developed for the intravenous treatment of COVID-19 in hospitalized patients and may be able to address different disease parameters within the one therapeutic treatment.

Brilacidin has been successfully tested in 8 clinical trials across multiple indications providing established safety and efficacy data on over 460 subjects. The drug has exhibited potent antibacterial activity in a Phase 2b trial in Acute Bacterial Skin and Skin Structure Infections (ABSSSI) and anti-inflammatory activity, as supported in Phase 2 clinical trials in Ulcerative Proctitis/Ulcerative Proctosigmoiditis and Oral Mucositis (see *Supplemental Information*). Brilacidin, through modulation of cyclic adenosine monophosphate (cAMP)/cyclic guanosine monophosphate (cGMP) pathway, is postulated to regulate the immune response based largely on its observed inhibition of phosphodiesterases (PDE4 and PDE3). Interestingly, numerous PDE inhibitors have been proposed as potentially promising anti-SARS-CoV-2 treatments by suppressing the heightened inflammatory response.**^94^**

Additional formulation work is planned for the inhaled delivery of brilacidin for prophylactic use, toward controlling infection in the nasal passage and lungs by leveraging brilacidin’s ability to inhibit SARS-CoV-2 by disrupting viral integrity and impacting viral entry. Such development efforts, if successful, may enable brilacidin to emerge as a particularly effective and differentiated antiviral by preventing and/or decreasing early infectivity due to SARS-CoV-2.

In this manuscript, we demonstrate brilacidin exhibits robust inhibition of SARS-CoV-2 in Vero cells and Calu-3 cells, supporting brilacidin as a promising novel drug candidate for the treatment of COVID-19. Functioning as a viral entry inhibitor,**^95, 96, 97^** proposed mechanisms of action for brilacidin include affecting the integrity of the viral membrane and preventing viral binding to cells. More detailed mechanistic studies are planned. Destabilizing viral integrity is a desirable antiviral property, especially in relation to pan-coronavirus agents, as the viral membrane is highly conserved and similar in construct across different coronavirus strains.**^98, 99^** Moreover, drugs that can directly disrupt viral integrity would be less prone to resistance due to mutation, unlike many antivirals, antibody-based treatments and vaccines that are currently in use and in development. Brilacidin exhibits an excellent synergistic inhibitory profile against SARS-CoV-2 in combination with remdesivir. Experiments conducted in endemic human coronaviruses are ongoing, with additional testing planned in other lethal coronaviruses (MERS-CoV, SARS-CoV), toward assessing the potential of brilacidin as a broad spectrum inhibitor of coronaviruses.

## Author Contributions

### Funding

George Mason University, with identified lead researcher Aarthi Narayanan, received financial support from Innovation Pharmaceuticals Inc. to conduct research on brilacidin’s antiviral properties.

## Acknowledgments

Authors thank William F. DeGrado (Department of Pharmaceutical Chemistry, University of California San Francisco) and Jun Wang (Department of Pharmacology and Toxicology, University of Arizona) for helpful discussions during this research project, and for manuscript review. Authors acknowledge contribution of Scott Van Wart from Enhanced Pharmacodynamics (ePD) to the analyses contributing to and generation of Figure 4. Authors thank IBT Bioservices for kindly providing the rVSV SARS-CoV-2 pseudovirus.

## Supplemental Information

Brilacidin has exhibited anti-inflammatory and antibacterial properties, clinically and preclinically: a summary is provided in Supplemental Information to this manuscript.

## Materials and Methods

### Cell Culture

Vero African green monkey kidney cells (ATCC, CCL-81), and Calu-3 human lung epithelial cells (ATCC, HTB-55), were obtained from the American Type Culture Collection. Vero cells were cultured in Dulbecco’s Modified Eagle’s Medium (DMEM, Quality Biological, 112-013-101CS) supplemented with 4.5g/L glucose, 2mM L-glutamine (FisherSci, MT2005CI), 5% heat-inactivated fetal bovine essence (FBE) (VWR, 10805-184) for Vero cells, 10ug/mL streptomycin and 10U/mL penicillin (VWR, 45000-652). Calu-3 cells were cultured in Eagle’s Minimum Essential Medium (EMEM, VWR, 670086) supplemented with 10% fetal bovine serum (FBS) (ThermoFisher, 10437028). All cell lines were cultivated at 37°C and 5% CO2.

### Inhibitors

Brilacidin (as brilacidin tetrahydrochloride) was provided by Innovation Pharmaceuticals Inc., and dissolved in dimethyl sulfoxide (DMSO) (Fisher Scientific, BP231). Hydroxychloroquine sulfate (SelleckChem, S4430), remdesivir (MedChemExpress, HY-104077), and favipiravir (FisherScientific, NC1312443) were obtained and dissolved in DMSO.

### Toxicity Screens

Cells were seeded in 96-well white plates 24h prior as follows: Vero and Caco-2 cells at 5×104 cells per well, Calu-3 cells at 1.3×105 cells per well. Inhibitors were diluted to the desired micromolar concentration (μM) in the appropriate culture media. Diluted compounds were added to the cells and plates were incubated at 37°C, 5% CO2. At 24 hours post-treatment (hpt), culture media was removed from the cells and cell viability was measured using a CellTiter-Glo^®^ Luminescent Cell Viability Assay per manufacturer’s instructions (Promega, G7572). Luminescence was measured using a Beckman Coulter DTX 880 multimode plate reader.

### SARS-CoV-2 Infections

SARS-CoV-2 (Washington strain 2019-nCoV/USA-WA1/2020) was obtained from BEI Resources (NR-52281) and was used for all infections, unless otherwise specified. For all infections, cells were seeded in 96-well plates 24h prior as follows: Vero cells at 5×104 cells per well, Calu-3 cells at 1.3×105 cells per well. Inhibitors were dissolved in DMSO and diluted in culture media to the indicated concentrations such that the final concentration of DMSO in the treatment was ≤0.1%. Mock infected cells were included as untreated and uninfected controls during all infections. Cells were pretreated with media containing drug or 0.1% DMSO vehicle control for 2h prior to infection. For non-direct viral infections, virus was diluted in culture media to the indicated multiplicity of infection (MOI) and this inoculum overlaid on cells for 1h. For direct viral infections, virus was diluted to the indicated MOI in culture media containing 0.1% DMSO or the inhibitor at the indicated concentration, and this virus:inhibitor solution, i.e., treated inoculum, incubated at 37°C and 5% CO2 for 1h. After this incubation, the treated inoculum was overlaid on cells for 1h. Conditioned media containing inhibitor or standard media was added to cells after removal of virus. For synergy experiments, fresh media containing inhibitor(s) was added to cells after removal of virus. Plates were incubated at 37°C, 5% CO2 for the indicated duration. At the indicated hour time point post-infection (hpi), viral supernatants were collected and stored at −80°C or used immediately for assays.

### Plaque Assay

Vero cells were plated in 12-well plates at a density of 2×105 per well and incubated for 24h. Infection supernatants were serially diluted to 10-4 in culture media and overlaid on cells for 1h. Cells were covered with Eagle’s Minimum Essential Medium (without phenol red, supplemented with 5% FBE, non-essential amino acids, 1mM sodium pyruvate (VWR, 45000-710), 2mM L-glutamine, 20U/mL penicillin, and 20μg/mL streptomycin) with 0.6% agarose (ThermoFisher, 16500100). At 48hpi, cells were fixed with 10% formaldehyde (FisherSci, F79P-4) for 1h. Medium was removed, wells were washed with diH2O, and stained with a 1% crystal violet (FisherSci, C581-25) and 20% ethanol solution (FisherSci, BP2818-4). Plaque assay datasets are represented as both plaque forming units per milliliter (PFU/mL) and as percentage of virus titer versus the DMSO control.

### Pseudovirus Spike Neutralization Assay

Vero cells were seeded in a black 96-well plate at 5×104 cells per well and incubated for 24h. Inhibitors were diluted in serum-free culture media at 2X of the indicated concentration and mixed with an equal volume of a luciferase-expressing pseudotyped virus (rVSV) containing the SARS-CoV-2 spike protein, diluted 1:5 in serum-free culture media (kindly contributed by IBT Bioservices) and incubated for 1h at room temperature. Media was removed from Vero cells and the rVSV:inhibitor solution was added to the cells and incubated for 1h at 37°C and 5% CO2. After 1h, equal volume of complete media was added to the cells and incubated for 24h at 37°C and 5% CO2. Media was removed and Vero cells were lysed with 1X Passive Lysis Buffer (Promega, E1941) for 30min while shaking. Equal volume of Luciferase Assay Substrate (Promega, E4550) was added to the cells and luciferase signal was measured using an integration time of 1s on a Beckman Coulter DTX 880 multimode plate reader. Neutralization was determined relative to the rVSV-only condition and mock-treated controls established limits of detection for relative light units.

### Confocal Micrcoscopy of Spike Neutralization Assay

Glass chamber slides (Ibidi, 80827) were treated with fibronectin from bovine plasma (Sigma-Aldrich, F1141) and allowed to dry, followed by washing with 1X phosphate buffered saline (PBS) (VWR, L0119). Vero cells were seeded in the fibronectin coated chamber slides at 5×104 cells per well 24h prior to assay. Cells were treated with rVSV:inhibitor solution in the same manner as previously described for the neutralization assay. At 1hpi and 4hpi, cells were fixed with 4% paraformaldehyde in 1X PBS (Alfa Aesar, J61899) for at least 10min followed by three washes with 1X PBS. Cells were permeabilized with 0.1% Triton X-100 (Sigma-Aldrich, X100) in 1X PBS for 15min at room temperature, followed by three washes with 1X PBS. Wells were blocked with 2% bovine serum albumin (BSA) (Fisher Scientific, BP1600) in 1X PBS while rocking overnight at 4°C. Cells were stained with SARS-CoV-2 Spike Antibody (ProSci, 3525) diluted to 1μg/mL in 0.1% BSA in 1X PBS and incubated at room temperature for 2h followed by three 5min washes with 1X PBS. Alexa Fluor™ 488 fluorescent dye-labeled secondary antibody (Invitrogen, A21206) was diluted 1:1000 in 0.1% BSA in 1X PBS and incubated on the chamber wells for 45min at room temperature protected from light, followed by three 5min washes with 1X PBS with 0.1% Tween-20 (Sigma-Aldrich, P9416). Mounting medium (Ibidi, 50001) with NucBlu™ DNA nuclear counterstain (Invitrogen, R37605) was added to all chamber wells and slides were protected from light prior to imaging. Mock-infected wells and primary antibody treatment only, or secondary antibody treatment only were included as assay and imaging controls, respectively. Images were obtained using a Nikon Eclipse Ti2 Fluorescent Microscope equipped with NIS Elements Software at 60X oil immersion objective. DAPI and FITC filters were utilized to qualitatively view fluorescence and quantitatively measure fluorescent counts of FITC using surface intensity plots. Neutralization was determined relative to the rVSV-only condition and mock-treated controls established background fluorescence.

### RNA extraction and RT-PCR

At the indicated time points post-infection, cells were washed with 1X PBS and lysed with TRIzol Reagent (Invitrogen). Intracellular RNA was extracted using Direct-zol Miniprep RNA kit (Zymo Research, R2051s) per manufacturer’s instructions. Extracted viral RNA was stored at −80°C or used immediately for analysis by RT-PCR. Primers and probes for detection of SARS-CoV-2 RNA specific to ORF 1ab for the envelope (E) and nucleocapsid (N) viral genome fragments were obtained from Integrated DNA Technologies (10006821, 10006822, 10006823). The probe was double-quenched with ZEN/IBFQ and contained a 6-FAM fluorescent dye attachment at the 5’ end. 18S rRNA endogenous control primer/probe set was utilized for semi-quantitative RT-PCR normalization (ThermoFisher, 4333760T). Thermal cycling conditions were adapted from Verso 1-step RT-qPCR kit (ThermoFisher, AB4101C) per the manufacturer’s instructions: 1 cycle at 50°C for 20min, 1cycle at 95°C for 15min, 40 cycles at 95°C for 15s with 52°C (CoV2 E,N) and 60°C (18S) for 1min using StepOnePlus™ Real Time PCR System. Controls with no template RNA and samples from mock infected controls were included for all analyses and established the limits of detection. Quantitative values were calculating using the ΔΔCt method with viral entities normalized to 18S levels and fold-changes calculated versus mock-infected conditions. Datasets are represented as both raw values calculated for fold-change and as a percentage versus the DMSO condition.

### Statistical analyses

Graphs represent the mean ± SD for all data obtained, with the exception of **Figure 4A** and **Figure 4B** for which sigmoidal Hill-type models as a function of brilacidin tetrahydrochloride concentration were fit to the data (using non-linear least squares regression in NONMEM Version 7.4, as performed by Enhanced Pharmacodynamics (ePD) on behalf of Innovation Pharmaceuticals Inc).

Statistical analyses and significance was determined using One-Way ANOVA with Dunnett’s Post Test in Prism 7 (Graph Pad) unless otherwise stated. Significance values are indicated using asterisks for *p<0.0332, **p<0.0021, ***p<0.0002, ****p<0.0001, ns for not significant.

