## Supplemental Information for "Brilacidin, a COVID-19 Drug Candidate, Exhibits Potent *In Vitro* Antiviral Activity Against SARS-CoV-2"

### A. Brilacidin Anti-Inflammatory Properties

**Figure S-A1. Brilacidin exhibited anti-inflammatory properties in a Phase 2 clinical trial in Ulcerative Proctosigmoiditis/Ulcerative Proctitis (UP/UPS).**

Clinical remission (with Endoscopic response) achieved after 6 weeks of treatment in >50% subjects in each cohort (50 mg, 100 mg, and 200 mg brilacidin, as daily retention enema)

Reduction in Inflammatory Biomarkers (by Cohort): Colonic Tissue Biopsies at Week 6 (Day 42)

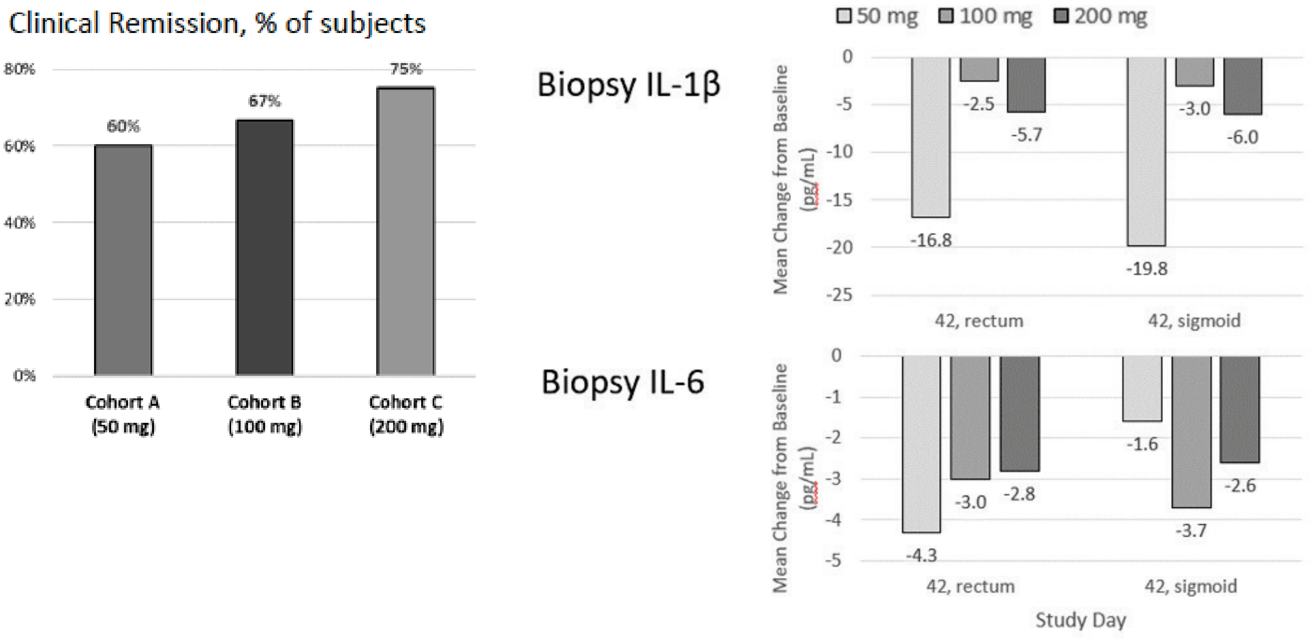

**Figure S-A2. Brilacidin exhibited anti-inflammatory properties in a Phase 2 clinical trial for attenuation of Severe Oral Mucositis (SOM) in patients with Head and Neck Cancer receiving chemoradiation.**

Brilacidin *oral rinse* demonstrated strongest therapeutic benefit in those Head and Neck Cancer patients on a 21-day (q3wk) cisplatin regimen

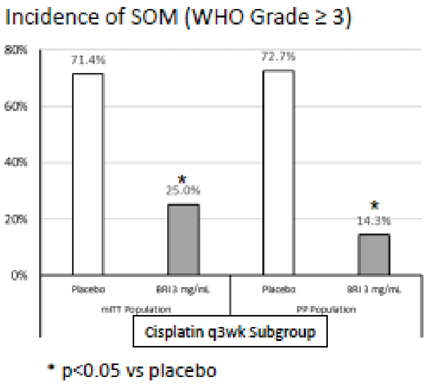


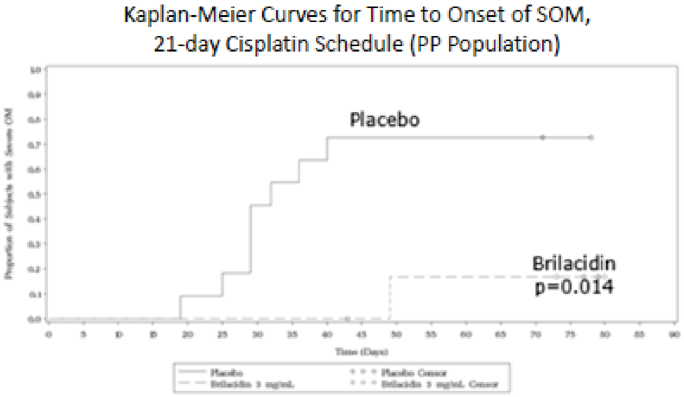


Note: period from approximately **19-49 days** during which SOM incidence rises strikingly in Placebo while not in the Brilacidin group

**Figure S-A3. Brilacidin exhibited anti-inflammatory properties in an *in vivo* mouse colitis model.**

In mice, dextran sulfate sodium (DSS) solution (5%) was administered to induce colitis, from Day 0 to Day 11

On Days 7-11, Brilacidin 400 mg/kg or 5-ASA 50 mg/kg or water (controls) were administered as a solution, per rectum

**Brilacidin** demonstrated **significant decrease by Day 11** compared to DSS-treated control gp for **Rectal bleeding** (by Hemoccult® kit), and **Stool consistency** also significantly more firm; results similar to positive control group (5-ASA)

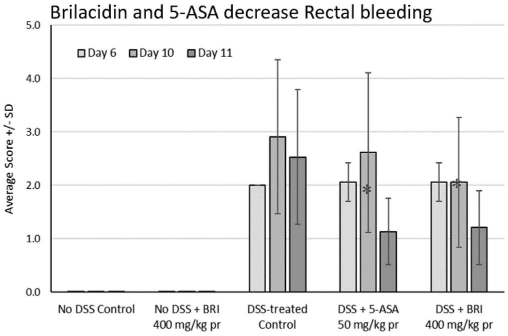


**Brilacidin** demonstrated **inhibition of IL-6 and IL-1β in distal colon tissue**

- IL-6 mean reduction of 40% in Brilacidin group, and of 86% in 5-ASA group, compared to DSS-treated control group
- IL-1β mean reduction of 27% in Brilacidin gp, and of 33% in the 5-ASA gp, compared to DSS-treated control gp

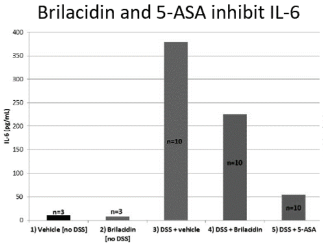


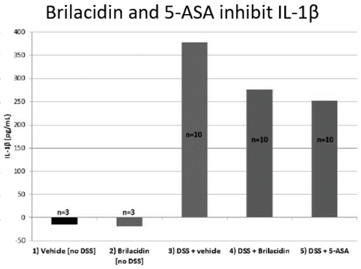

**Table S-A1. Brilacidin inhibits multiple pro-inflammatory cytokines and chemokines *in vitro* and *ex vivo.***

| Study Type | Test System | IPI Study Number | Brilacidin $IC_{50}$ | Unit |
|---|---|---|---|---|
| Inhibition of human PDE4B2 | in vitro PDE-Glo™ PDE assay | 2014-05-29 | 3 | µM |
| Inhibition of human PDE3A | in vitro PDE-Glo™ PDE assay | 2014-07-18 | 1.8 | µM |
| Inhibition of LPS-induced TNF-α release | Rat alveolar macrophage (NR8383) cells; ELISA assay | 2014-08-21 | 442 | nM |
| Inhibition of LPS-induced MMP-9 release | Rat alveolar macrophage (NR8383) cells; ELISA assay | 2014-11-10 | 2.3 | µM |
| Inhibition of LPS-induced MCP-1 release | Rat alveolar macrophage (NR8383) cells; ELISA assay | 2014-11-13 | 750 | nM |
| Inhibition of LPS-induced IL-6 release | Rat alveolar macrophage (NR8383) cells; ELISA assay | 2014-12-03 | 274 | nM |
| Inhibition of LPS-induced IL-1β release | Rat alveolar macrophage (NR8383) cells; ELISA assay | 2015-02-10 | 702 | nM |
| Inhibition of LPS-induced CINC-3 release | Rat alveolar macrophage (NR8383) cells; ELISA assay | 2015-02-16 | 425 | nM |
| Inhibition of LPS-induced TNF-α release | Human monocytic leukemia (THP-1) cells; ELISA assay | 2016-07-21 | 23.4 | µM |
| Inhibition of LPS-induced IL-8 release | Human monocytic leukemia (THP-1) cells; ELISA assay | 2016-08-02 | 10.8 | µM |

Source: IPI Research Reports. Reports present only one experiment dataset, although multiple experiments with similar findings were archived.
CINC = cytokine-induced neutrophil chemoattractant; ELISA = enzyme-linked immunosorbent assay; IL = interleukin; LPS = lipopolysaccharide; MCP = monocyte chemoattractant protein; MMP = matrix metalloproteinase; PDE = phosphodiesterase; TNF = tumor necrosis factor

Methods/Summary for Table S-A1.

Using the *in vitro* PDE-Glo™ phosphodiesterase assay, brilacidin was demonstrated to inhibit human PDE4B2 enzyme and human PDE3A enzyme in a dose-dependent manner with an $IC_{50}$ of approximately 3 µM and 1.8 µM, respectively. PDE4 and PDE3 inhibition results in subsequent down-regulation of pro-inflammatory cytokines/chemokines and its regulators, and experiments were conducted to confirm such down-regulation by brilacidin in *ex vivo* cell-based assays. In these assays, cells were exposed to brilacidin for 45 minutes before an 8-hour lipopolysaccharide (LPS) stimulation in the presence of brilacidin; cytokine/enzyme concentrations were determined by ELISA using a standard immunoassay kit. Results demonstrated dose-dependent down-regulation of the LPS-response within treated cells, as would be expected from reduction in activation of the nuclear factor-κB (NF-κB) pathway (stimulated by LPS bound to cell membrane toll-like receptors) from PDE inhibition by brilacidin.

### B. Brilacidin Antibacterial Properties

**Figure S-B1. Brilacidin exhibited antibacterial properties in two phase 2 trials in Acute Bacterial Skin and Skin Structure Infections (ABSSSI); Brilacidin efficacy compared favorably to Daptomycin.**

Efficacy by Primary Endpoint: Early Clinical Response

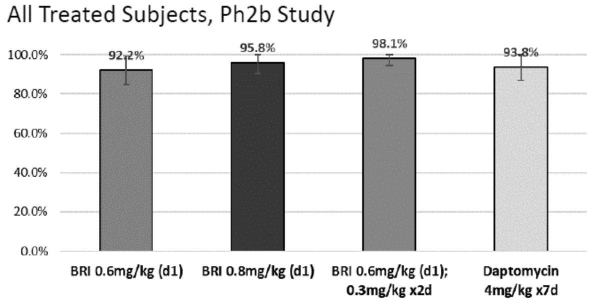


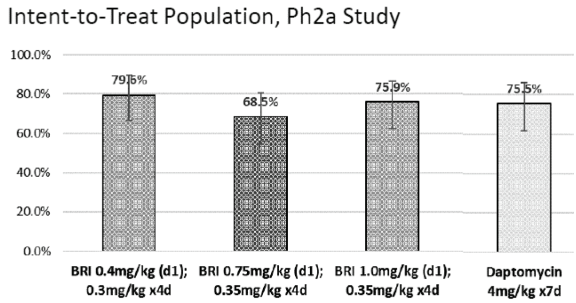


BRI: Brilacidin

**Table S-B1. Investigator Assessment: Clinical Success Rates by Baseline Pathogen (Phase 2b study).**

| Baseline Pathogen | PI Clinical Assessment at Day 7/8: EOT | | | | PI Clinical Assessment at Day 10-14: STFU | | | |
|---|---|---|---|---|---|---|---|---|
| | Brilacidin | | | Daptomycin | Brilacidin | | | Daptomycin |
| | 0.6<br>1 day | 0.8<br>1 day | 0.6/0.3<br>3 days | 7 days | 0.6<br>1 day | 0.8<br>1 day | 0.6/0.3<br>3 days | 7 days |
| *Staphylococcus aureus* | | | | | | | | |
| MSSA only | 16/17 (94.1) | 15/18 (83.3) | 12/13 (92.3) | 11/13 (84.6) | 16/17 (94.1) | 14/17 (82.4) | 12/12 (100.0) | 11/12 (91.7) |
| + *S. lugdunensis* | 1/1 (100.0) | | 1/1 (100.0) | | 1/1 (100.0) | | 1/1 (100.0) | |
| + *S. anginonsus-milleri* | | 1/1 (100.0) | 1/1 (100.0) | | | 1/1 (100.0) | 1/1 (100.0) | |
| + *S. pyogenes* | | | | 2/2 (100.0) | | | | 2/2 (100.0) |
| MRSA only | 9/9 (100.0) | 7/8 (87.5) | 10/11 (90.9) | 12/13 (92.3) | 9/9 (100.0) | 6/7 (85.7) | 8.8 (100.0) | 11/12 (91.7) |
| + *E. faecalis* | | | | 1/1 (100.0) | | | | 1/1 (100.0) |
| + *S. agalactiae* | | | | 1/1 (100.0) | | | | 1/1 (100.0) |
| *Streptococcus* | | | | | | | | |
| *agalactiae* | | | | 1/1 (100.0) | | | | 1/1 (100.0) |
| *anginonsus-milleri* | 2/2 (100.0) | 2/3 (66.7) | 2/3 (66.7) | 3/3 (100.0) | 2/2 (100.0) | 2/3 (66.7) | 2/3 (66.7) | 3/3 (100.0) |
| *pyogenes* | 1/1 (100.0) | | | | 1/1 (100.0) | | | |
| *Staphylococcus* | | | | | | | | |
| *lugdunensis* | | 1/1 (100.0) | | 1/1 (100.0) | | 1/1 (100.0) | | 1/1 (100.0) |
| *Enterococcus* | | | | | | | | |
| *faecalis* | | | | 1/1 (100.0) | | | | 1/1 (100.0) |
| *Group C Beta-hemoloytic streptococci* | | | | 2/2 (100.0) | | | | 2/2 (100.0) |

Blank cells = no participants presented with the specified baseline pathogen.

EOT: End of Treatment. STFU: Short-Term Follow-Up.

**Table S-B2. Brilacidin shows potent broad-spectrum activity against gram-positive bacteria, with coverage against gram-negative bacteria.**

| **Gram-Positive** | | **MIC (ug/ml)* 2-3 isolates/organism** | | |
|---|---|---|---|---|
| | **Brilacidin** | *Linezolid* | *Vancomycin* | *Ceftazidime* |
| Entero. faecalis | 1 | 1-2 | 1 | >64 |
| Entero. faecium (VRE) | 1 | 1-2 | >128 | >64 |
| Staph. aureus (MRSA) | 0.5 - 1 | 1-2 | 0.5- 1 | 32 |
| Staph. epidermidis | 0.25 - 0.5 | 0.5 - 1 | 2 | 16-32 |
| Staph. saprophyticus | 0.25 - 0.5 | 1-2 | 1-2 | 32 - >64 |
| Staph. spp. (coagulase-) | 0.25 - 0.5 | 1 | 1-2 | 16-32 |
| Strept. agalactiae | 2 | 1 | 0.5 | 0.5 |
| Strept. pneumoniae | 4-8 | 1 | 0.5 | 0.25 |
| Strept. pyogenes | 1-4 | 1 | 0.5 | 0.12 |
| Strept. viridians | 2-8 | 1 | 0.5 - 1 | 0.5 - 4 |

| **Gram-Negative** | | **MIC (ug/ml)* 2-3 isolates/organism** | | |
|---|---|---|---|---|
| | **Brilacidin** | *Ceftazidime* | *Linezolid* | *Vancomycin* |
| Citrobacter fruendi | 2-4 | 0.25 - 2 | >16 | >128 |
| Citrobacter koseri | 1-2 | 0.12 - 0.25 | >16 | >128 |
| Enterobacter cloacae | 0.5 - 4 | 0.25 | >16 | >128 |
| Escherichia coli | 1-2 | 0.06 | >16 | >128 |
| Klebsiella oxtoca | 2-8 | 0.06 - 0.12 | >16 | >128 |
| Klebsiella pneumoniae | 1-2 | 0.06 - 0.12 | >16 | >128 |
| Morganella morganii | 2 - >64 | 2-16 | >16 | >128 |
| Proteus mirabilis | 64 - >64 | 0.03-0.06 | >16 | >128 |
| Proteus vulgaris | 64 - >64 | 0.12 | >16 | >128 |
| Providencia stuartii | 16-64 | 0.12 - 64 | >16 | >128 |
| Acinetobacter spp. | 4 | 2-64 | >16 | 128 - >128 |
| Pseud. aeruginosa | 32 | 1-8 | >16 | >128 |
| Serratia marcescens | 32 | 0.12 - 0.25 | >16 | >128 |
| Stenotrophomonas maltophilia | 8 - >64 | 4-8 | >16 | 32-128 |
| Haemophilus influenzae | 4-8 | 0.06 - 0.12 | 16 - >16 | 128 |

Methods: Broth microdilution assays performed according to standard CLSI guidelines.

**Table S-B3. Brilacidin shows activity against drug-susceptible and drug-resistant defined phenotypes in Staphylococcus species.**

| | *Drug-Susceptible* | *OXA-R* | *VRSA/VISA, OXA-R* | *LZD, OXA-R* | *DAP-NS, OXA-R* | *VRSA/VISA, DAP-NS, OXA-R* |
|---|---|---|---|---|---|---|
| **Brilacidin** MIC | 0.25-1 | 0.25-2 | 0.5-1 | 0.5-1 | 0.5-2 | 0.5-1 |
| # of isolates | 217 | 161 | 7 | 5 | 5 | 3 |

Acronyms: OXA-R: oxacillin-resistant; VRSA: vancomycin-resistant to S. aureus; VISA: vancomycin intermediate S. aureus; LZD-NS: linezolid non-susceptible; DAP-NS: daptomycin non-susceptible

Methods: Broth microdilution assays performed according to standard CLSI guidelines.

Summary: Brilacidin was active *in vitro* against all isolates of S. aureus and coagulase-negative staphylococci, including isolates of S. aureus with characterized resistance to daptomycin, linezolid, and vancomycin. Against S. aureus isolates, there was no alteration in activity against resistant isolates relative to susceptible isolates. Against coagulase-negative staphylococci, activity was not affected by resistance to methicillin.

**Figure S-B2. Brilacidin exhibits potent and rapid bactericidal activity (from 0.5 to 6 hours) against E. coli and S. aureus.**

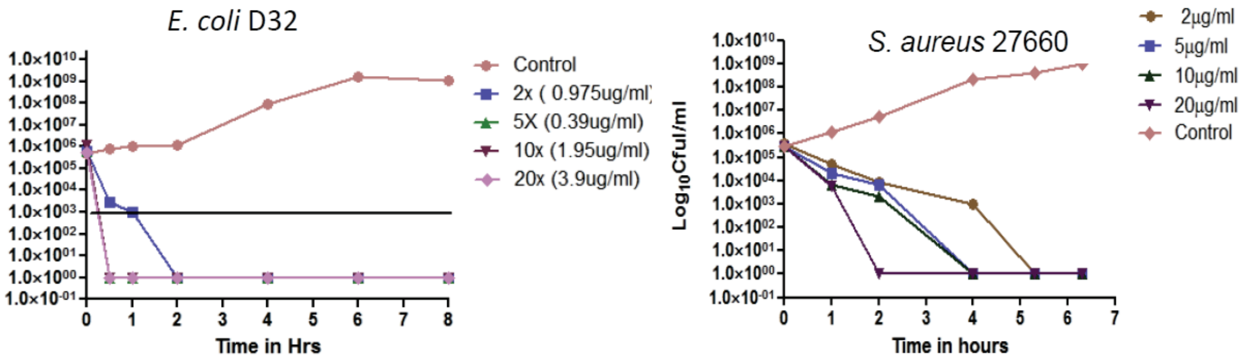


Time-kill a measure of CFU/ml after exposure of E. coli D32 and S. aureus 27660 to brilacidin.
Methods: Broth microdilution assays performed according to standard CLSI guidelines.

**Figure S-B3. Brilacidin exhibits potent and rapid bactericidal activity against stationary phase cultures of Methicillin-Susceptible Staph Aureus (MSSA) and Methicillin-Resistant Staph Aureus (MRSA).**

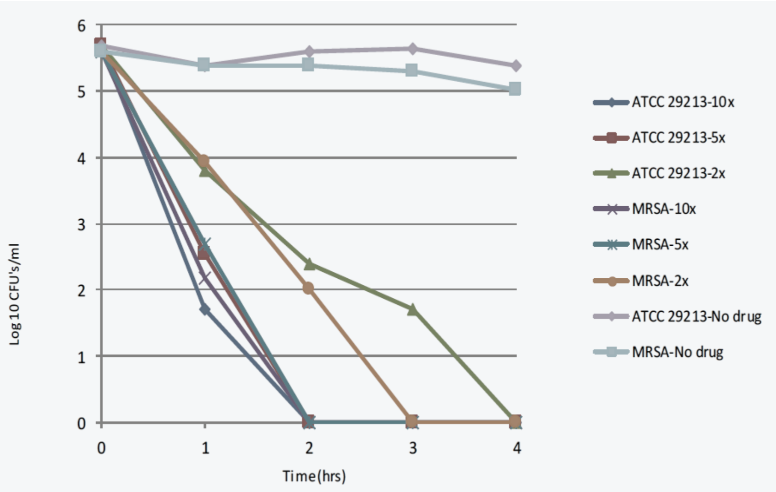


Methods: Broth microdilution assays performed according to standard CLSI guidelines.
Time-kill at >$10^3$ $Log_{10}$ Reductions of ≤ 2 hours at two 2x MIC.

Daptomycin showed little antimicrobial activity up to 10x the MIC (data not shown).

**Table S-B4. Brilacidin demonstrates potent inhibition and non-cytotoxic selectivity for bacteria over mammalian cells.**

| PMX compound | MIC or $MIC_{90}$* (µg/ml) | Cytotoxicity ($EC_{50}$ µg/ml) | | | Selectivity ($EC_{50}$/MIC) | | |
|---|---|---|---|---|---|---|---|
| | *S. aureus* | RBCs | 3T3 | HepG2 | RBCs | 3T3 | HepG2 |
| 30063 | 1.0* | >500 | 430 | 1,031 | 558 | 430 | 1,031 |
| melittin | 2 | 2 | 4 | 1 | 1 | 2 | 0.5 |
| HDPs | 2 - 5 | 20 - 50 | | | 10 - 20 | | |

Acronyms: Red Blood Cells (RBCs); Host Defense Proteins/Peptides (HDPs)

Methods: Cytotoxicity of brilacidin (PMX-30063) was evaluated using human erythrocytes by $OD_{414}$ measurements for hemoglobin concentration and using calorimetric assay in human liver cell line (HepG2) and an embryonic mouse cell line (NIH/3T3). This assay measures the bioreduction of a novel tetrazolium compound to a soluble formazan product by viable cells.

Summary: Brilacidin is smaller than natural HDPs (1/5$^{th}$ to 1/10$^{th}$) while exhibiting comparable or greater potency while 50 to 100 folds more selective.

**Figure S-B4. Brilacidin has low risk for bacterial resistance developing based on serial passage assays.**

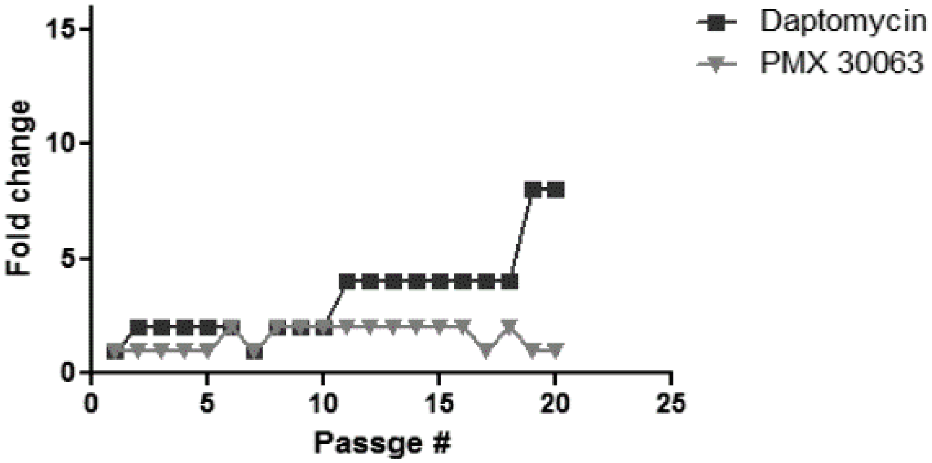


Methods: Broth microdilution assays performed according to standard CLSI guidelines.
